# Spatial gradient in activity within the insula reflects dissociable neural mechanisms underlying context-dependent advantageous and disadvantageous inequity aversion

**DOI:** 10.1101/243428

**Authors:** Xiaoxue Gao, Hongbo Yu, Ignacio Saez, Philip R. Blue, Lusha Zhu, Ming Hsu, Xiaolin Zhou

## Abstract

Humans are capable of integrating social contextual information into decision-making processes to adjust their attitudes towards inequity. This context-dependency emerges both when individual is better off (i.e. advantageous inequity) and worse off (i.e. disadvantageous inequity) than others. It is not clear however, whether the context-dependent processing of advantageous and disadvantageous inequity rely on dissociable or shared neural mechanisms. Here, by combining an interpersonal interactive game that gave rise to interpersonal guilt and different versions of the dictator games that enabled us to characterize individual weights on aversion to advantageous and disadvantageous inequity, we investigated the neural mechanisms underlying the two forms of inequity aversion in the interpersonal guilt context. In each round, participants played a dot-estimation task with an anonymous co-player. The co-players received pain stimulation with 50% probability when anyone responded incorrectly. At the end of each round, participants completed a dictator game, which determined payoffs of him/herself and the co-player. Both computational model-based and model-free analyses demonstrated that when inflicting pain upon co-players (i.e., the guilt context), participants cared more about advantageous inequity and became less sensitive to disadvantageous inequity, compared with other social contexts. The contextual effects on two forms of inequity aversion are uncorrelated with each other at the behavioral level. Neuroimaging results revealed that the context-dependent representation of inequity aversion exhibited a spatial gradient in activity within the insula, with anterior parts predominantly involved in the aversion to advantageous inequity and posterior parts predominantly involved in the aversion to disadvantageous inequity. The dissociable mechanisms underlying the two forms of inequity aversion are further supported by the involvement of right dorsolateral prefrontal cortex and dorsomedial prefrontal cortex in advantageous inequity processing, and the involvement of right amygdala and dorsal anterior cingulate cortex in disadvantageous inequity processing. These results extended our understanding of decision-making processes involving inequity and the social functions of inequity aversion.

## Introduction

Inequity aversion, or the preference for fairness, is a common other-regarding preference observed in human society (Bolton and Ockenfels, 2000; Brosnan and de Waal, 2014; Decety and Yoder, 2016; Fehr and Schmidt, 1999; Tricomi et al., 2010). When facing conflict between self-interest and norm of equity, the aversion to inequity motivates individuals to make decisions that are relatively equal such as sharing resources with others (Brosnan and de Waal, 2014; Decety and Yoder, 2016; Fehr and Schmidt, 1999), and rejecting or even punishing others’ unfair treatment (Dawes et al., 2007; Fehr and Fischbacher, 2003; Fehr and Gächter, 2002; Sanfey et al., 2003). Such motivations of inequity-aversion are thus fundamental for human large-scale cooperation (Brosnan and de Waal, 2014; Fehr and Schmidt, 1999) and social justice (Rawls, 1958; Reeve, 1998).

The aversion to inequity does not occur in social vacuum; in fact, we encounter inequity in various social contexts. Individuals are capable of integrating social context-related information into decision-making processes to adjust their responses to inequity aversion, which refers to the *context-dependency of inequity aversion* (Bäker et al., 2012; Güroğlu et al., 2010; Loewenstein et al., 1989; Tversky and Simonson, 1993; Wright et al., 2011). These flexible adjustments take place regularly in various contexts in our everyday life, such as the psycho-physiological states of the decision-maker (Harlé and Sanfey, 2007; Hu et al., 2015; Hu et al., 2014), the traits of the person with whom the individual interacts (Loewenstein et al., 1989; Radke et al., 2012; Wu et al., 2011), and the societal/economic environment (Dhami, 2016). This ability to make flexible adjustments enables us to maintain cooperative relationship, maximize self-interest, and adapt to dynamic social environments (John et al., 2013; Lenartowicz et al., 2010; Louie and De Martino, 2013; Rilling and Sanfey, 2011; Rushworth et al., 2007). Thus increased knowledge of the neural mechanisms underlying these adjustments can provide valuable clues into not only the understanding of various social behaviors but also the mechanisms underlying mental dysfunctions involving social maladaptation, such as autism (Palmer et al., 2015) or psychopathy (Domes et al., 2013).

Over the past few decades, a series of economic and neuroimaging studies on decision-making involving inequity aversion have led to the development of several classic paradigms (e.g. dictator game, DG^1^, and ultimatum game, UG^2^; (Kahneman et al., 1986)) and computational models (e.g. the Fehr-Schmidt inequity-aversion model; Fehr and Schmidt, 1999), which extend our understanding on the psychological and neural mechanisms underlying inequity aversion under decontextualized experimental environments (i.e. general inequity aversion; For reviews, see Aoki et al., 2015; Fehr and Camerer, 2007; Feng et al., 2015; Gabay et al., 2014). As for the context-dependency of inequity aversion, an abundance of studies (For review, see Wang et al., 2015) focused on how individuals perceive and respond to unfair offers (i.e. disadvantageous offers) as responders in UG across different social contexts; findings from these studies suggest that brain networks involved in fairness considerations are highly context-dependent and are dissociable from brain networks involved in fairness processing in decontextualized environment (Wright et al., 2011). Despite these advances, in most of these neuroimaging studies, inequity aversion is regarded as a general concept and the primary focus is on the situations where participants received less than the others (i.e. the disadvantageous inequity), such as the responder in UG. Little attention was paid to the situation where participants received more than the others (i.e. the advantageous inequity). However, as shown in behavioral studies, individual reactions to inequity may differ in terms of the relative status of the individual (i.e. advantageous status or disadvantageous status). Although the utility of an action decrease when both types of inequity increase, people’s responses to advantageous inequity is not as strong as to disadvantageous inequity (Dawes et al., 2007; Fehr and Schmidt, 1999; Loewenstein et al., 1989). Rejecting advantageous inequity to themselves requires more cognitive resources than rejecting disadvantageous inequity to themselves (Van den Bos et al., 2006). Consistently, it is shown that these two forms of inequity aversion emerge at different developmental (Blake et al., 2015; Blake and McAuliffe, 2011; Fehr et al., 2008; McAuliffe et al., 2013; McAuliffe et al., 2017) and evolutionary (Brosnan, 2009; Brosnan and de Waal, 2014) stages.

Of note, the aversions to advantageous and disadvantageous inequity vary differently across contexts. For example, when distributing resources as a dictator, individuals tend to avoid getting more when interacting with cooperative others (e.g. friends or neighbors), but are more tolerant of advantageous inequity when interacting with competitive others (e.g. competitors or salesmen). In contrast, this context manipulation has no effect on disadvantageous inequity aversion (Loewenstein et al., 1989). This behavioral dissociation naturally leads to the question as to whether there exist dissociable neural mechanisms underlying the context-dependent processing of advantageous and disadvantageous inequity aversion. The answer to this question is surprisingly unknown.

Answering this question will contribute to distinguish between two alternative hypotheses regarding the neural mechanisms for inequity processing in the brain (Figure 3B). One hypothesis (Hypothesis 1; Figure 3B-I) is that a common neural mechanism is adopted for encoding both advantageous and disadvantageous inequity, analogous to the common neural currency hypothesis in neuroeconomics which posits that reward value from various modalities is encoded by the same neural computational process (Levy and Glimcher, 2012; Sugrue et al., 2005). This shared inequity encoding system is then qualified by a separate system that represents advantageous versus disadvantageous status. Critically, this hypothesis posits that it is this latter status-encoding system that is modulated by the attributes of social interaction context. According to this account, if we can identify brain regions that show different sensitivity to inequity across contexts (i.e. the interaction effect between context and inequity level), we should observe *overlapping* brain regions that encodes the level of inequity common to advantageous and disadvantageous status and across different social interaction contexts, along with brain areas arbitrating the relative weights of the advantageous and disadvantageous status. The alternative hypothesis (Hypothesis 2; Figure 3B-II) is that advantageous inequity aversion and disadvantageous inequity aversion may be two separate psychological constructs with dissociable neural mechanisms (i.e. the dissociable inequity aversion hypothesis). In that case, we should observe that the manipulation of context modulate *non-overlapping* brain networks for advantageous and disadvantageous inequity. Distinguishing these two hypotheses could improve our understanding of the context-dependent adjustments of inequity aversion and its role in maintaining harmonious social interaction. To this end, we need an interaction context that can simultaneously modulate individuals’ advantageous and disadvantageous inequity aversion.

Here we combine an interpersonal interactive game (Yu et al., 2014a) and a variant of the DG (Saez et al., 2015), which enable us to measure inequity-related decision-making in a common social context, i.e. the interpersonal guilt context. Interpersonal guilt is one of the most common social/moral emotions in daily life. It has been shown that guilt is closely related to inequity aversion because the former both signals and constitutes the obligation of wrong-doers to balance the inequity created by their wrong-doing or transgression (Baumeister et al., 1994; Owens, 2008; Tangney et al., 2007). On the one hand, obtaining more or suffering less than the others are important sources of guilt (Baumeister et al., 1994; Hassebrauck, 1986; Homans, 1961; Kubany and Watson, 2003; Walster et al., 1978). In DG, the extent to which a dictator is averse to advantageous inequity is regarded in some decision theory on fairness as reflecting anticipatory guilt (Fehr and Schmidt, 1999; Krajbich et al., 2009; Rey-Biel, 2008). On the other hand, the experience of guilt motivates prosocial behaviors, such as making compensations, which are aimed at removing inequity and restoring the relationship back to an even footing (Baumeister et al., 1994; Izard, 1977). Consistent with this notion, previous studies have showen that individuals are willing to compensate the victim when guilt is induced, indicating their increased generosity to the victims (De Hooge et al., 2007; Katelaar and Au, 2003; Nelissen and Dijker, 2007; Reed, 2010; Yu et al., 2014a). This increased generosity may result from increased advantageous inequity aversion and/or decreased disadvantageous inequity aversion when making decisions in a state of guilt (Saez et al., 2015). We first tested this hypothesis in two behavioral studies. Participants played a dot-estimation task with anonymous co-players who received pain stimulation with 50% probability when either of the two co-players responded incorrectly. At the end of each round, participants completed a continuous version (cf. Saez et al., 2015; Experiment 1) or a binary choice version of DG (Experiment 2) as the dictator, which determined the payoffs of him/herself and co-players. The weights on advantageous and disadvantageous inequity aversion during monetary allocation under different conditions were estimated using the Fehr-Schmidt inequity aversion model (Fehr and Schmidt, 1999; Saez et al., 2015), in which individuals trade off between self-interest and two forms of inequity aversion. Our behavioral results suggested that the context manipulation could simultaneously modulate individuals’ advantageous and disadvantageous inequity aversion, and the contextual effects on two forms of inequity aversion were uncorrelated with each other at the behavioral level.

Based on the behavioral results, we further investigate whether there are dissociable neural mechanism underlying context-dependent advantageous and disadvantageous inequity aversion. Participants performed Experiment 3 (the same paradigm as Experiment 2) in an MRI scanner. Neuroimaging results revealed that the context-dependent processing of advantageous inequity aversion involved the left anterior insula (aINS), the right dorsolateral prefrontal cortex (DLPFC), and dorsomedial prefrontal cortex (DMPFC), while the context-dependent processing of disadvantageous inequity aversion involved the left posterior insula (pINS), the amygdala, and dorsal anterior cingulate cortex (dACC). The representation of inequity aversion exhibited a spatial gradient in activity within the insula, such that anterior parts predominantly involved in advantageous inequity aversion processing and posterior parts predominantly involved in disadvantageous inequity aversion processing.

## Methods

### Participants

#### Experiment 1 (behavioral study)

Thirty-seven healthy graduate and undergraduate students took part in Experiment 1. Three participants were excluded from data analysis because of failing to responses in more than 10 trials (1 participant) or misunderstanding instructions (2 participants), leaving 34 participants (23 females; mean age 20.0 years; age range: 18–23 years) for data analysis.

#### Experiment 2 (behavioral study)

Twenty-eight healthy graduate and undergraduate students took part in Experiment 2. Three participants with low accuracy in the catch trials (1 participant) or failing to respond in more than 10 trials (2 participant) were excluded from the following analysis, leaving 25 participants (15 females, mean age 20.3 years; age range: 18–25 years) for data analysis.

#### Experiment 3 (fMRI study)

Thirty-four right-handed healthy graduate and undergraduate students took part in the fMRI scanning. Eight participants with excessive head movements (>3mm, 5 participants), low accuracy in the catch trials (2 participants) or failing to respond in more than 10 trials (1 participant) were excluded from the following analysis, leaving 26 participants (17 females, mean age 20.1 years; age range: 18–24 years) for data analysis. One participant showed no unique solution for individual-level model fitting and was excluded from analyses involving individual-level model fitting.

None of the participants in all the three experiments reported any history of psychiatric, neurological, or cognitive disorders. Informed written consent was obtained from each participant before each experiment. All the studies were carried out in accordance with the Declaration of Helsinki and were approved by the Ethics Committee of the School of Psychological and Cognitive Sciences, Peking University.

### Procedure

#### Experiment 1 (behavioral study; Continuous version of DG)

Three same sex participants came to the experiment room together. Upon arrival they met three same sex confederates and were told that they would later play an interactive game together through an intranet in separate rooms. The interactive game was composed of 3 A players and 3 B players and the participants were always peusdorandomly assigned to be A players. An intra-epidermal needle electrode was attached to the left wrist of each participant for cutaneous electrical stimulation (Inui et al., 2002). Each participant experienced several pain stimulations and rated each of them on a scale of 1 (“no feeling”) to 8 (“unbearable”). The final intensity of pain stimulation was calibrated to a subjective pain rating of “6”, which was a moderate punishment for the participants. The participants were told that the pain stimulation that each B player would receive in the interactive game would be the one that each B player rated as a 6 in pain intensity, which was also moderate punishment for them.

The experiment consisted of 2 phases. In the first phase, each participant performed a pre-experiment task in which they performed as the dictator in DG and made 20 monetary decisions (see *Continuous version of DG and choice setting;* Figure 1B). In each trial, the participant was randomly paired with one of the anonymous co-players and decided the self-payoff and payoff of this co-player by indicating the tokens he/she wanted to give to the co-player. The participant was told that their co-players were performing other tasks at the same time, and the payoffs of the participant and co-players in this phase depended on the participant’s decisions. After the experiment, one of the decisions made by the participant in this phase was selected randomly, which determined the final payoffs of the participant and the co-player (1 point for 0.2 yuan). Each participant’s performance in this task was regarded as baseline condition in the following data analysis.

**Figure 1.**
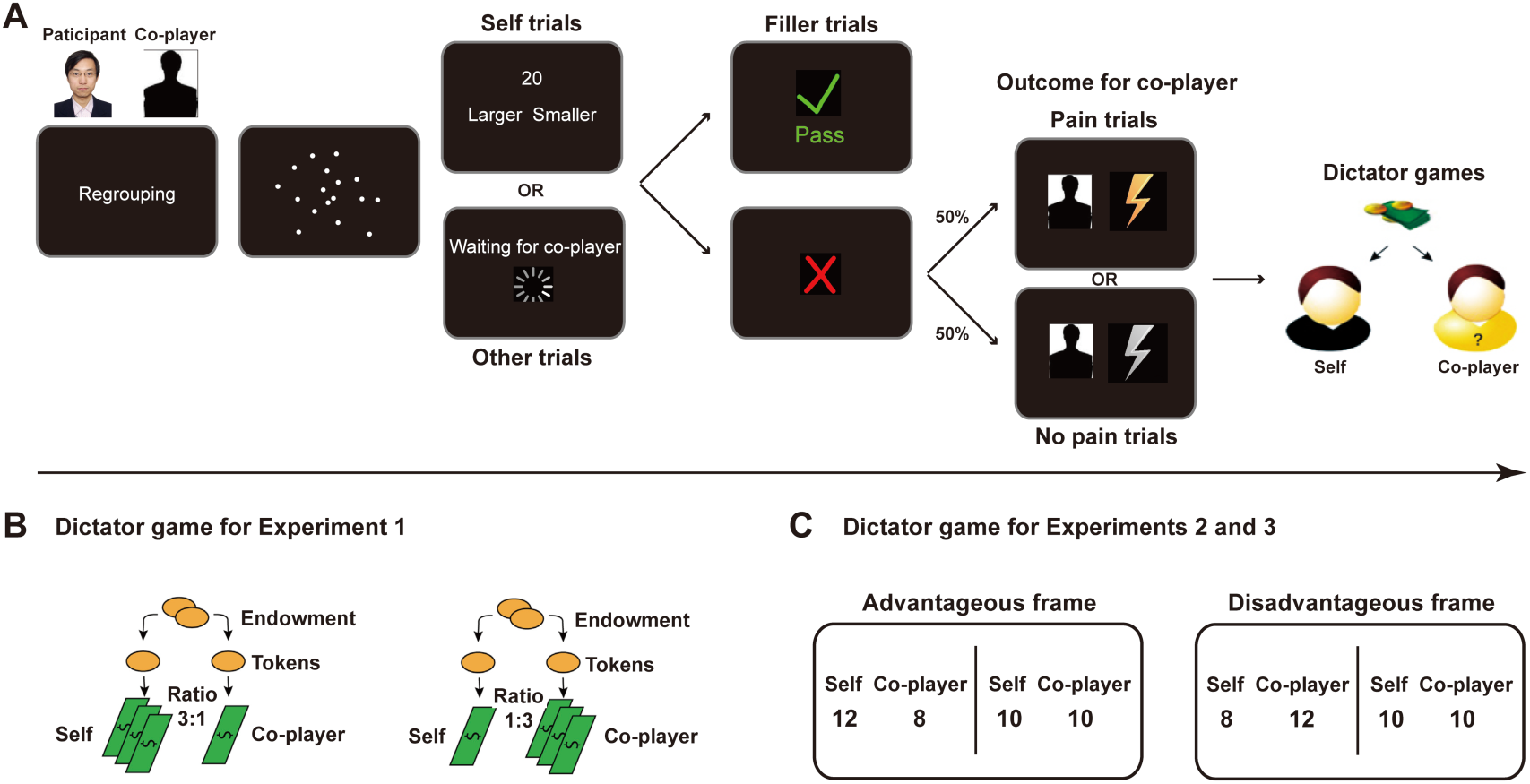
Task displays, timing, and design. **(A)** Each trial began by informing the participant that the program had randomly chosen one out of the three B players as his/her co-player in the current trial, the identity of whom was unknown. Sometimes the participant had to quickly estimate the number of dots on the screen and sometimes he/she needed to wait for his/her co-player to make the estimation. The participant was told to press a corresponding button, using the left or right thumb, to indicate whether his/her estimation was larger or smaller than the number (randomly chosen from 19, 20, and 21) appearing on the next screen. There were always 20 dots on the screen, the positions of which were randomly generated. The positions of ‘Smaller’ and ‘Larger’ were counterbalanced across participants. If the answer was correct, no one would accept the pain stimulation, and the game entered the next trial. If anyone’s answer was incorrect, the co-player in the current trial had a 50% chance of being administered the pain stimulation, which was randomly decided by the computer. If the co-player needed to receive the pain stimulation, a yellow flash was presented; if the co-player did not need to receive the pain stimulation, a gray flash was presented. Then at the end of each estimated incorrect trials, the participant acted as Dictator in the dictator game to determine the payoff for him/herself and the co-player. **(B)** Two versions of dictator games were used. In Experiment 1 (left), in each choice, the participant was faced with an endowment of T tokens and could unilaterally choose to give some tokens to the co-player. The relative cost and benefit of giving were manipulated by independently varying how much each token was worth to the participant and the co-player. For example, a ratio of 1 to 3 meant that 1 token was worth 1 point for the participant and 3 point for the co-player, and vice versa (Saez et al., 2015). Given that in the DG from Experiment 1, the advantageous status and disadvantageous status were determined by participants’ choices, which would result in an unbalanced design matrix in the fMRI analysis, we further developed a binary choice version of DG in Experiments 2 and 3 (right). Each binary choice consisted of two options representing the payoff given to the participant and the co-player after the experiment. One option was always “10 points for me, and 10 points for the co-player” and the other option was an unequal option with different values in each trial. To dissociate advantageous and disadvantageous status, there were two types of unequal option which corresponding to the two kinds of status. No matter what version the DG is, one trial was selected after the experiment to determine the final payoff of the participant and the co-player in this trial. In the fMRI experiment, before and after the screen showing dot estimation outcome and whether the co-player whould receive the pain stimulation and before and after the presentation of each binary choice, a fixation cross (not shown in the figure) was presented for a variable of interval ranging from 1 to 5s. These were for the purpose of fMRI signal deconvolution.

Subsequently, in the second phase, participants performed the interactive task described in Figure 1A with 3 co-players. Each trial began by informing the participant that the program had randomly chosen one out of the three B players as his/her co-player in the current trial, the identity of whom was unknown. Sometimes the participant had to quickly estimate the number of dots on the screen, and sometimes he/she needed to wait for his/her co-player to do the estimation. If the answer was correct, no one would accept the pain stimulation, and the current trial terminated. If anyone’s answer was incorrect, the co-player in the current trial had 50% chance to accept the pain stimulation, which was randomly decided by the computer. Then the participant should perform as dictator in DG and made a monetary choice at the end of each estimating-incorrect trial to decide self-payoff and payoff of the co-player in this trial (see *Continuous version of DG and choice setting*; Figure 1B). After the experiment, one decision that the participant made was randomly selected from all of their decisions, which determined the payoffs for the participant and the co-player in this task (1 point for 0.2 yuan).

The correct trials (i.e. the participant estimates correctly or the co-player estimates correctly) were treated as filer trials (40 trials, half of these trials were participant-estimating trials). There were four possible conditions for the incorrect trials (the participant estimated incorrectly or the co-player estimated incorrectly) forming a 2 (Agent: Self vs. Other) × 2 (Outcome: Pain vs. Nopain) within-subject design: the participant estimated incorrectly and the co-player received pain stimulation (Self_Pain), the participant estimated incorrectly and the co-player did not receive pain stimulation (Self_Nopain), the co-player estimated incorrectly and the co-player received pain stimulation (Other_Pain), the co-player estimated incorrectly and the co-player did not receive pain stimulation (Other_Nopain). Given that the regret of estimating incorrectly and the empathy for co-players’ pain could be two confounding factors here, we introduced Nopain (Self_Nopain and Other Nopain) conditions to control for the regret of estimating incorrectly and the Other_Pain condition to control for the empathy for the co-players’ pain. Thus, in the current study, the interaction effect between Agent and Outcome were the effect that we focused on (i.e. guilt effect). The experiment consisted of 120 trials (20 trials for each of the above four conditions and 40 correct trials). Each condition consisted of 20 monetary choices (1 for each trial, see *Continuous version of DG and choice setting),* which appeared randomly in the end of each condition.

#### Experiment 2 (behavioral study; binary-choice version of DG)

The role assignment and pain rating procedure in Experiment 2 was the same as in Experiment 1. The main interactive task was similar to Experiment 1, with the exception that the participant made four sequential monetary binary choices (see *Binary choice dictator game and choice setting* for details; Figure 1C) between an equal option (i.e. standard option) and an unequal option (i.e. test option) at the end of each incorrect trial. The increased number of binary choices was to increase the quality of both model fitting and fMRI analysis. Each option in the binary choice represented the payoff that the participant and the co-player in this trial would earn. After the experiment, one binary choice was selected randomly from the all the decisions that the participant made, and this determined the payoffs for the participant and the co-player in this task (1 point for 1 yuan).

The condition setting was the same as Experiment 1. Experiment 2 consisted of 108 trials (18 trials for each of the above four conditions and 36 correct trials, half of these trials were participant-estimating trials). Each condition consisted of 72 monetary binary choices (four for each trial, see *Binary-choice version of DG and choice setting;* Figure 1C), which appeared randomly in the end of each condition.

#### Experiment 3 (fMRI study; binary-choice version of DG)

Each participant came to the scanning room individually. Upon arrival he/she met three same sex confederates and was told that they would later play an interactive game together through intranet, but in separate rooms. The interactive game was comprised of one A player and three B players and the participants were always the A players.

The pain rating procedure, task, and condition setting in the fMRI study were the same as in Experiment 2. The scanning session consisted of three runs and lasted for ~57 min. Each run consisted of 36 trials (six trials for each of the above four conditions and twelve correct filler trials).

For all of the three experiments, after the main interactive task, each participant rated their feeling of guilt corresponding to the four conditions on a scale of 1 (‘not at all’) to 7 (‘very strong’).

### Dictator games

#### Continuous version of DG and choice setting

In Experiment 1, in the pre-experiment task and at the end of each estimating-incorrect trial in the interactive task (Figure 1A), the participant made a monetary choice at the end of each estimating-incorrect trial, which was a continuous version of DG (Figure 1B) with an expanded choice space that allowed us to dissociate the behavioral effects of advantageous inequity aversion and disadvantageous inequity aversion (Saez et al., 2015). In each choice, the participant was faced with an endowment consisting of T tokens and could unilaterally choose to give some tokens (To) to the co-player in this trial. The relative cost and benefit of giving in each trial was manipulated by independently varying how much each token was worth to the participant (rs) and the co-player (ro). For example, rs equaled 1 and ro equaled 3 meant that 1 token was worth 1 point (0.2 yuan) for the participant but 3 point for the co-player, and vice versa. Table S1 listed 20 combinations of To, ro and rs used in the experiment according to Saez et al. (2015), which were the same for the baseline condition in the pre-experiment task and each of the four conditions in the interactive task in Experiment 1and appeared randomly in each condition. This paradigm also enabled us to use the Fehr-Schmidt inequity aversion model (Fehr and Schmidt, 1999; Saez et al., 2015) to dissociate the weight of advantageous inequity aversion and disadvantageous inequity aversion in decision-making at the group level.

#### Binary-choice version of DG and choice setting

Given that in the continuous version of DG of Experiment 1, the advantageous status (i.e. participants allocated more to themselves than co-players) and disadvantageous status (i.e. participants allocated more to co-players than themselves) were determined by participants’ choices, thus resulting in an unbalanced design matrix for fMRI analysis, in Experiment 2 and 3, we further developed a binary choice version of DG to dissociate individual advantageous and disadvantageous inequity aversion at both the behavioral level and the neural levels (Figure 1C). At the end of each estimating-incorrect trial, the participant made four sequential monetary binary choices. Each binary choice consisted of two options representing the payoff given to the participant (Ms) and the payoff given to the co-player (Mo) after the experiment if the participant chose this option and this choice was chosen. One option was always “10 points for me, and 10 points for the co-player” (i.e. the standard option) and the other option was unequal option, which was one of the 72 test options in Table S2. Two frames of test options, the advantageous frame (e.g., “18 points for me, and 12 points for the co-player”) and the disadvantageous frame (e.g., “12 points for me, and 18 points for the co-player”), were implemented to dissociate advantageous inequity aversion and disadvantageous inequity aversion. The binary choice sets were the same for the four conditions. The abstract difference of inequity between each option pair was not correlated with the abstract difference of self-payoff *(R* = -0.057, *p* = 0.634) or the abstract difference of other-payoff (*R* = 0.127, *p* = 0.288) between them.

Notably, in all the three experiments, several instructions were applied to exclude potential confounding factors. First, to control for the effect of reputation, participants were told that their co-players did not know they had the right to make allocation choices and could not see their choices during the experiment. Second, to exclude the possibility that participants tried to counter-balance the payoff of the three co-players, the entire interpersonal game was anonymous, and participants did not know any information regarding the co-player in each trial. Thus, it was better for all the co-players to make decisions according to the particular context in that particular trial. Third, to exclude the possibility that participants tried to counter-balance self-payoff and other-payoff across 4 binary choices in the end of each trial (Experiments 2 and 3), they were informed that the binary choices were independent from each other and the numbers in each binary choice were randomly selected by the computer. Thus, the best strategy for them was to make every decision independently, because they could not expect how the options would change in the subsequent binary choices.

#### Computational Modeling

**Fehr-Schmidt inequity aversion model (Model 1)**. We used the Fehr-Schmidt inequity aversion model (Fehr and Schmidt, 1999; Saez et al., 2015) to dissociate the weight of advantageous inequity aversion and disadvantageous inequity aversion in decision-making. Specifically, we defined the subjective value function as:

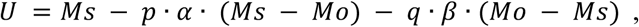

where Ms and Mo refers to self and other payoff respectively, and p and q are indicator functions: p =1, if Ms ≥ Mo (advantageous inequity) and 0 otherwise; and q = 1, if Ms < Mo (disadvantageous inequity), and 0 otherwise. Thus, α and β quantify subjective aversion to inequity under advantageous and disadvantageous frames, respectively.

The function was calibrated to choice behavior by using a softmax specification with inverse temperature parameter λ, such that in each trial, the probability of the participant choosing one option is given by

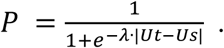

In all the three experiments, to test the effect of guilt context on inequity aversion, we conducted maximal likelihood estimation at group level by maximizing the log likelihood function over individual participant i and trial t:

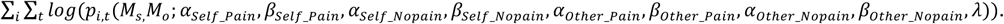

The SEs of estimated parameters were obtained through a bootstrap procedure with 200 iterations.

In Experiments 2 and 3, to test whether the contextual effects on advantageous and disadvantageous inequity aversion correlated with each other, we further conducted maximal likelihood estimation at the individual level by maximizing the log likelihood function over each participant i and trial t with 1000 different starting values:

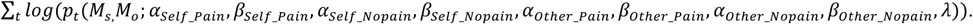

The individual parameters generated from the maximum log likelihood estimation were accepted for further analysis. We further compared the Fehr-Schmidt inequity aversion model (Fehr and Schmidt, 1999; Saez et al., 2015) with 7 alternative models in Experiments 2 and 3. See *Supplementary Material* for more information.

#### Behavioral measures of individual sensitivity to contexts

To test whether individual sensitivity to contexts predicted individual context-dependent neural responses to advantageous inequity aversion and disadvantageous inequity aversion, every participant was asked to complete the Guilt And Shame Proneness scale (GASP). The GASP was applied to measure individual differences in the proneness to experience guilt and shame across a range of personal transgressions (Cohen et al., 2011). This questionnaire contains two guilt subscales that assess negative behavior-evaluations (guilt-NBE) and repair action tendencies (guilt-repair) following private transgressions and two shame subscales that assess negative self-evaluations (shame-NSEs) and withdrawal action tendencies (shame-withdraw) following publically exposed transgressions. People with high guilt-NBE scores feel guiltier after harming others and are more empathic, humble, conscientious, agreeable, and altruistic than those with low guilt-NBE scores. Thus, the scores of guilt-NBE in GASP were used as index for individual sensitivity to interpersonal guilt in daily life in the data analysis.

#### Data acquisition and preprocessing

Images were acquired using a GE Healthcare 3.0 T Medical Systems Discovery MR 750 with a standard head coil at Peking University (Beijing, China). T2*-weighted echoplanar images (EPI) were obtained with blood oxygenation level-dependent (BOLD) contrast. Thirty-five transverse slices of 4 mm thickness that covered the whole brain were acquired in an interleaved order (repetition time = 2000 ms, echo time = 30 ms, field of view = 192×192 mm^2^, flip angle = 90°).

The fMRI data were preprocessed and analyzed using Statistical Parametric Mapping software SPM8 (Wellcome Trust Department of Cognitive Neurology, London). Images were slice-time corrected, motion corrected, resampled to 3 mm × 3 mm × 3 mm isotropic voxels, normalized to MNI space, spatially smoothed with an 8 mm FWHM Gaussian filter, and temporally filtered using a high-pass filter with a cutoff frequency of 1/128 Hz.

#### FMRI data analysis

**Whole-brain analysis.** According to our fMRI hypothesis (Figure 3A), we performed event-related fMRI analyses of participants’ neural responses on the decision phase at which they decided self payoffs and other payoffs. To address the question as to whether there were dissociable brain systems for context-dependent advantageous inequity and disadvantageous inequity processing, choices were classified into advantageous frame and disadvantageous frame according to the inequity type of test options (i.e. unequal options), and each choice in each frame was further medium split into high-inequity choices (HI) and low-inequity choices (LI) (Table S2). Thus, both the advantageous frame and the disadvantageous frame consisted of 8 conditions: 2 (Agent: Self vs. Other) × 2 (Outcome: Pain vs. Nopain) × 2 (Inequity level: HI vs. LI).

The first general linear model (GLM1) was specified to address our main question as to whether there exist and, if so, what are the dissociable neural mechanisms underlying context-dependent advantageous and disadvantageous inequity aversion. Separate regressors in GLM1 were specified for fMRI responses to:

- R1-R8: The presentation of options in each of the 8 conditions for advantageous frame, i.e. Self_Pain_LI, Self_Pain_HI, Self_Nopain_LI, Self_Nopain_HI, Other_Pain_LI, Other_Nopain_HI, Other_Nopain_LI, and Other_Nopain_HI;
- R9-R16: The presentation of options in each of the 8 conditions for disadvantageous frame, i.e. Self_Pain_LI, Self_Pain_HI, Self_Nopain_LI, Self_Nopain_HI, Other_Pain_LI, Other_Nopain_HI, Other_Nopain_LI, and Other_Nopain_HI;
- R17-R20: The revelation of incorrect outcomes for 4 outcome conditions, i.e. Outcome_Self_Pain, Outcome_Self_Nopain, Outcome_Other_Pain, and Outcome_Other_Nopain;
- R21: Dot estimation screen, which consisted of the presentation of dots and estimation response in the trials that participants estimated the number of dots;
- R22: Waiting screen, which consisted of the presentation of dots and waiting period for co-player’s estimation in the trials that the co-player estimated the number of dots;
- R23: The revelation of correct outcomes;
- R24: The presentation of equal options;
- R25: Missed allocation, i.e. the period for missed decisions;
- R26 – R31: Six movement parameters, which were treated as regressors of no interest.

In the first-level (within-participant) statistical analysis, for each of the 4 main conditions (i.e. Self_Pain, Self_Nopain, Other_Pain, Other_Nopain) in the advantageous frame, we defined a main effect contrast corresponding to the difference between the neural responses to high inequity choices and low inequity choices: HI > LI. Then in the second-level (group-level) analysis, these four contrast maps for each participant were fed into a flexible factorial design. We defined an interaction effect contrast corresponding to the effect of the guilt context on the weight of advantageous inequity aversion: (Self_Pain - Self_Nopain > Other_Pain - Other_Nopain) (Table S5). Similar analyses were conducted for disadvantageous frame to model the effect of guilt context on the weight of disadvantageous inequity aversion (Table S6). All the results were corrected for multiple comparisons using voxel-wise threshold of p<0.001 (uncorrected) combined with cluster-level threshold of p<0.05 (23 voxels, FWE- corrected). This cluster-level threshold (number of voxels in the cluster) was determined using a Monte Carlo simulation (Ledberg et al., 1998) as implemented in the AFNI AlphaSim program (http://afni.nimh.nih.gov/pub/dist/doc/manual/AlphaSim.pdf). To further test whether there exist overlapping context-dependent processing for two form of inequity aversion, two further analyses were conducted: (1) conjunction analysis was conducted using the contrast maps for advantageous and disadvantageous frames (Table S5 and S6); (2) we combine the data of two frames to conduct contrast analysis together.

We established two more GLMs to exclude the possible confounding effect of the value representation of self-payoff (GLM2) and other-payoff (GLM3). Since we hypothesized the same neural responses to self-payoff in advantageous and disadvantageous frames and the same neural responses to other-payoff in these two frames, we combined the fMRI data of advantageous and disadvantageous frames in GLM2 and GLM3. In GLM2 choices in each of the 4 main conditions were medium split into high self-payoff options (HS) and low self-payoff (LS) according to self-payoff in the test options. Thus, the first 8 regressors in GLM2 were specified for fMRI responses to the presentation of options in each of the 8 conditions i.e. Self_Pain_LS, Self_Pain_HS, Self_Nopain_LS, Self_Nopain_HS, Other_Pain_LS, Other_Nopain_HS, Other_Nopain_LS, and Other_Nopain_HS. Similarly, for GLM3, choices in each of the 4 main conditions were medium split into high other-payoff options (HO) and low other-payoff (LO) according to other-payoff in the test options. Thus, the first 8 regressors in GLM2 were specified for fMRI responses to the presentation of options in each of the 8 conditions i.e. Self_Pain_LO, Self_Pain_HO, Self_Nopain_LO, Self_Nopain_HO, Other_Pain_LO, Other_Nopain_HO, Other_Nopain_LO, and Other_Nopain_HO. The other regressors for GLM2 and GLM3 were the same as R17-R31 in GLM1. Contrasts for GLM2 and GLM3 were set according to Table S9 and Table S10, respectively.

## PPI analysis

To test the probability that the processing of context-dependent advantageous and disadvantageous inequity aversion relied on not only on distinct neural activations but also distinct functional connectivity, we performed a psychophysiological interaction analysis (PPI; Friston et al., 1997) focusing on the left aINS, rDLPFC, and DMPFC identified in the advantageous frame and the left pINS identified in the disadvantageous frame (see *Results).* In the advantageous frame, the left aINS, rDLPFC and DMPFC seeds were created with spheres of radius 3 mm centered on peak MNI coordinates of the three regions in the whole-brain analysis (left aINS [-30, 21, -20], rDLPFC [39, 20, 37], DMPFC [-12, 47, 40], Figure 4A). In the advantageous frame, for each seed, we conducted a PPI analysis for each of the 4 main conditions (i.e. Self_Pain, Self_Nopain, Other_Pain, Other_Nopain) to assess differential functional connectivity with this seed in HI condition compared with LI condition. For each PPI analysis, the BOLD signal within the seed was used as the physiological factor and the HI versus LI contrast as the psychological factor. At the individual level, each GLM was specified with one regressor representing the extracted time series in the seed (i.e. the physiological variable), one regressor representing the psychological variable of interest (i.e. HI > LI), and a third regressor representing the interaction of the preceding two regressors (the PPI term). This GLM was estimated to reveal areas whose activation was predicted by the PPI term, with the physiological and the psychological regressors being controlled. At the group-level, for each seed, four beta maps representing the PPI effect in the four main conditions for each participant was fed into a flexible factorial design. We defined an interaction effect contrast corresponding to the effect of the guilt context on the weight of advantageous inequity aversion: (Self_Pain - Self_Nopain > Other_Pain - Other_Nopain) (Table S5). Similar PPI analysis was conducted for the disadvantageous frame to access differential functional connectivity with the left pINS [-42, -4, 7] (Figure 4B). We defined an interaction effect contrast corresponding to the effect of guilt context on the weight of disadvantageous inequity aversion: (Self_Nopain - Self_Pain > Other_Nopain - Other_Pain) (Table S6). Small volume correction (SVC) (Worsley et al., 1996) was used for multiple comparisons applied at P_FWE_ < 0.05, following an initial threshold of *P* < 0.005 (uncorrected). ROIs were identified as functional brain regions implicated in emotion processing (i.e. bilateral amygdala), conflict processing (i.e. dACC), reward processing (i.e. bilateral ventral striatum (VS) and ventrolmedial prefrontal cortex (VMPFC)) (For reviews, see Aoki et al., 2015; Fehr and Camerer, 2007; Feng et al., 2015; Gabay et al., 2014) were identified as ROIs. The peak MNI coordinates of these regions were extracted from activation maps generated from meta-analyses on “emotion”, “conflict” and “reward” terms using the Neurosynth database (http://neurosynth.org; Yarkoni et al., 2011), which were independent from the current study. For SVC, the small volumes were defined as spheres with 10mm radius, centered on the peak MNI coordinates of these regions: left amygdala [-20, -6, -18], right amygdala [26, -4, -20], dACC [6, 28, 36], left VS [-12, 10, -8], right VS [12, 10, -8], VMPFC [2, 48, -14].

## ROI analysis

### Gradient effect

We quantified spatial gradients in advantageous and disadvantageous inequity aversion processing within the insula by extracting parameter estimates from five anatomical ROIs (3 voxels × 3 voxels × 3 voxels cubes; [-30, 20, -17], [-33, 14, -11], [-39, 8, -5], [-40, 2, 1], [-42, -4, 7]) along the axis from the left aINS identified in the advantageous frame to the left pINS identified in the disadvantageous frame. To exclude the regional correlations derived from smoothing, we re-estimated the GLM1 without smoothing in preprocessing and conducted the ROI analysis. For each ROI, we extracted the beta estimates for each condition in GLM1, evaluated the absolute value of the 3-way interaction effects [(Self_Pain_High > Self_Pain_Low) - (Self_Nopain_High > Self_Nopain_Low) - (Other_Nopain_High > Other_Pain_Low) + (Other_Nopain_High > Other_Nopain_Low)] for the advantageous frame and disadvantageous frame, and then entered them into a 2 (context: Advantageous vs. Disadvantageous) × 5 (ROI locations) repeated-measures one-way ANOVA.

### Individual difference analysis

To test whether neural mechanisms underlying individual differences in contextual effects on advantageous and disadvantageous inequity aversion were dissociable from each other, we conducted ROI analyses for brain regions involved in advantageous inequity aversion processing (i.e. left aINS, rDLPFC, and DMPFC) and brain regions involved in disadvantageous inequity aversion processing (i.e. left pINS, dACC, and right Amygdala). For each ROI, we extracted the beta estimates for each condition for each participant, and evaluated the value of 3-way interaction effects [(Self_Pain_High > Self_Pain_Low) - (Self_Nopain_High > Self_Nopain_Low) − (Other_Nopain_High > Other Pain_Low) + (Other_Nopain_High > Other_Nopain_Low)] for the advantageous frame and disadvantageous frame respectively, corresponding to the strength of neural adjustments in these regions (the greater the difference between this value and zero, the stronger the neural adjustments are). Then we examined whether individual sensitivity to contexts (guilt-NBE score in GASP) predicted these interaction effects using liner regression.

### Mapping the locations of the reported insular clusters in previous studies using UG on insula

Many studies have investigated the neural mechanisms under decision involving disadvantageous inequity under different contexts using UG (in which participants acted as responders) and have suggested the important role of insula in these decisions (Table S12). To closely examine the roles of different insular sub-regions in these studies, we parceled the insular clusters into the ventral anterior, dorsal anterior, and mid-posterior clusters by applying the insular sub-regions template (k = 3 solutions; Kelly et al., 2012), and then mapped the MNI coordinates of insula identified in previous studies using UG on these 3 sub-regions (Figure S7). To dissociate the role of insula in general disadvantageous inequity aversion (regardless of contexts) and context-dependent disadvantageous inequity aversion, the results of unfair offers > fair offers (regardless of contexts) contrast and context-related contrasts were mapped in different colors. This map showed that most of the insular regions identified in the unfair offers > fair offers contrast are located on the dorsal anterior insula, while the insular regions identified in the context-related contrasts were located on both the dorsal anterior and the mid-posterior parts of insula.

## Results

### Dissociable Contextual Effects on Advantageous and Disadvantageous Inequity at Behavioral Level

For all three experiments, 2 (Agent: Self vs. Other) × 2 (Outcome: Pain vs. Nopain) repeated measures analyses of variance (ANOVA) on the self-reported guilt ratings in the post-scan questionnaire yielded significant interaction effects between Agent and Outcome (Table S3; Figure 2D). Compared with Nopain conditions, inflicting pain upon the co-player increased the Agent effect on guilt, which demonstrated the robustness and validity of our paradigm to induce guilt.

**Figure 2.**
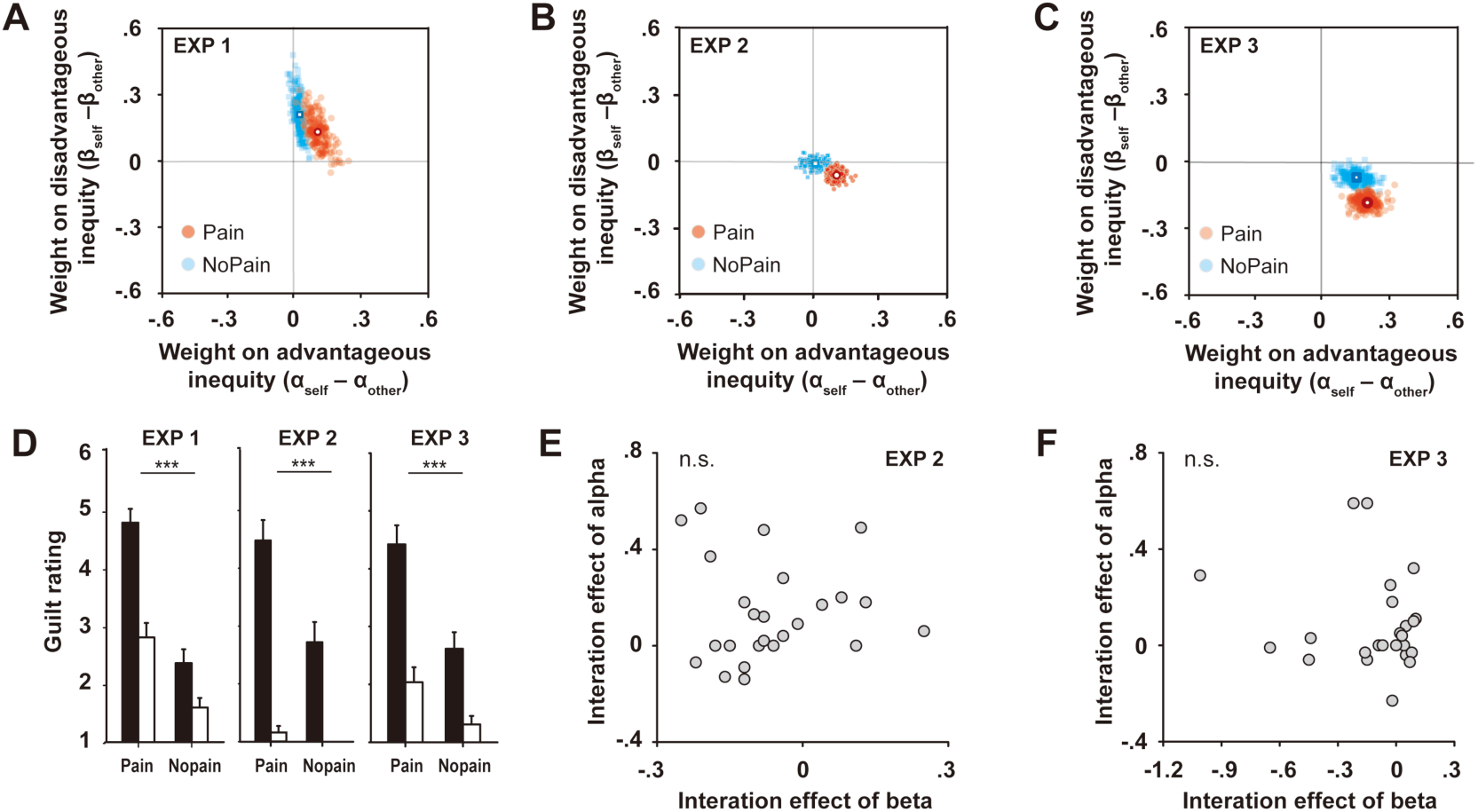
Behavioral results. **(A - C)** Group-level model-based results in Experiments 1, 2, and 3, respectively. The x axis and y axis represent the difference between Self condition and Other condition for the weight on advantageous inequity (α) and the weight on disadvantageous inequity (*β*), respectively. The red dots represent bootstrap pseudo-sample estimates (see *Computational Modeling)* for Pain condition, while the blue dots represent bootstrap pseudo-sample estimates for Nopain condition. Thus, the relative location of red dots to blue dots captures the interaction effect between Agent and Outcome (i.e. the guilt effect) on α and β. In all three experiments, guilt induced increased *α* and decreased *β* (red dots move down-right relative to the blue dots), indicating participants’ increased sensitivity to advantageous inequity but decreased sensitivity to disadvantageous inequity. **(D)** Guilt rating in the four conditions in all three experiments, respectively. **(E and F)** Individual-level model-based results in Experiments 2 and 3, respectively. Each figure represents the relationship between the interaction effect between Agent × Outcome on *α* and the interaction effect between Agent × Outcome on *β*. Our results showed that the interaction effect on α and β were uncorrelated with each other, which means the person with higher contextual effect on alpha does not necessarily have the same effect for beta.

First, to test whether our context manipulation modulated individual advantageous inequity and disadvantageous inequity aversion, we used the Fehr-Schmidt inequity aversion model (Fehr and Schmidt, 1999; Saez et al., 2015) to capture individual weight on advantageous inequity aversion and weight on disadvantageous inequity aversion in decision-making (see *Computational Modeling*). Both group-level model fitting in all the three experiments and individual level model fitting in Experiments 2 and 3 demonstrated that compared with Nopain conditions, causing pain to others significantly increased the agent effect on α and decreased the agent effect on β (Figure 2A-C and Figure S2; Table S3 and S4). These results indicated that our context manipulation could simultaneously modulate individuals’ advantageous and disadvantageous inequity aversion. Consistent patterns of model-free results and model-based results were observed for all DGs in the three experiments (see *Supplementary Material).* Statistical descriptions for model-free results and model-based results were listed in Table S3 and S4, respectively.

Second, to test whether the contextual effects on advantageous and disadvantageous inequity are dissociable at the behavioral level, we examined the relationship between the contextual effect on α and β (i.e. the interaction effects between Agent and Outcome on α and β) in both Experiments 2 and 3. Our results showed that the contextual effects on α and β were uncorrelated with each other (Experiment 2: *r* = .023, *p* = .911; Experiment 3: *r* = .152, *p* = .479), which means that the person with higher contextual effect on *α* does not necessarily have the same effect for *β* (Figure 2E-F). In sum, our behavioral results demonstrated the modulation effect of guilt context on inequity aversion and suggested that the contextual effects on two forms of inequity aversion are dissociable at the behavioral level.

The Fehr-Schmidt inequity aversion model explained participants’ choices significantly better than the other 7 plausible alternative models (see *Model comparison* and *Table S11* in *Supplementary Material* for more information).

### Dissociable Neural Responses to Inequity in Advantageous and Disadvantageous Frames

Our behavioral results demonstrated that when feeling guilty, participants’ weight on advantageous inequity aversion (parameter α) increased and their weight on disadvantageous inequity aversion (parameter *β*) decreased. These contextual effects on advantageous and disadvantageous inequity aversion are dissociable at the behavioral level. In order to address the question as to whether there are dissociable brain systems for advantageous inequity processing and disadvantageous inequity processing, binary choices were classified into the advantageous frame and disadvantageous frame, and each type of choices was further medium split into high-inequity choices (HI) and low-inequity choices (LI). In close correspondence to the behavioral analysis, we established the fMRI hypotheses showing the potential response pattern for brain regions that involved in context-dependent advantageous and disadvantageous inequity aversion processing (Figure 3A). Specially, given the increased advantageous inequity aversion induced by guilt, we hypothesized that brain regions representing context-dependent advantageous inequity aversion would show increased sensitivity to advantageous inequity in the Self_Pain condition, which would in turn result in boosted activity differences between HI and LI conditions in these regions in advantageous frame (Figure 3A-I). Thus, the activity of these brain regions would show significant Agent × Outcome × Inequity level 3-way interaction effects [(Self_Pain_High > Self_Pain_Low) − (Self_Nopain_High > Self_Nopain_Low) − (Other_Nopain_High > Other_Pain_Low) + (Other_Nopain_High > Other_Nopain_Low)] in the advantageous frame. Similarly, brain regions representing context-dependent disadvantageous inequity aversion would show reduced sensitivity to disadvantageous inequity in the Self_Pain condition, which would in turn result in decreased activity difference between HI and LI conditions in these regions in the disadvantageous frame (Figure 3A-II). Thus, the activity of these regions would also show significant Agent × Outcome × Inequity level 3-way interaction effects, but the direction of this effect would be opposite to that in the advantageous frame [(Self_Pain_Low > Self_Pain_High) − (Self_Nopain_ Low > Self_Nopain_ High) − (Other_Nopain_ Low > Other_Pain_ High) + (Other_Nopain_ Low > Other_Nopain_ High)]. Based on these hypotheses, we focused on the neural responses on the decision phase at which participants decided self and other payoffs and established GLM1 to reveal brain regions that were involved in context-dependent processing of advantageous inequity aversion and disadvantageous inequity aversion (see *FMRI data analysis* for details).

**Figure 3.**
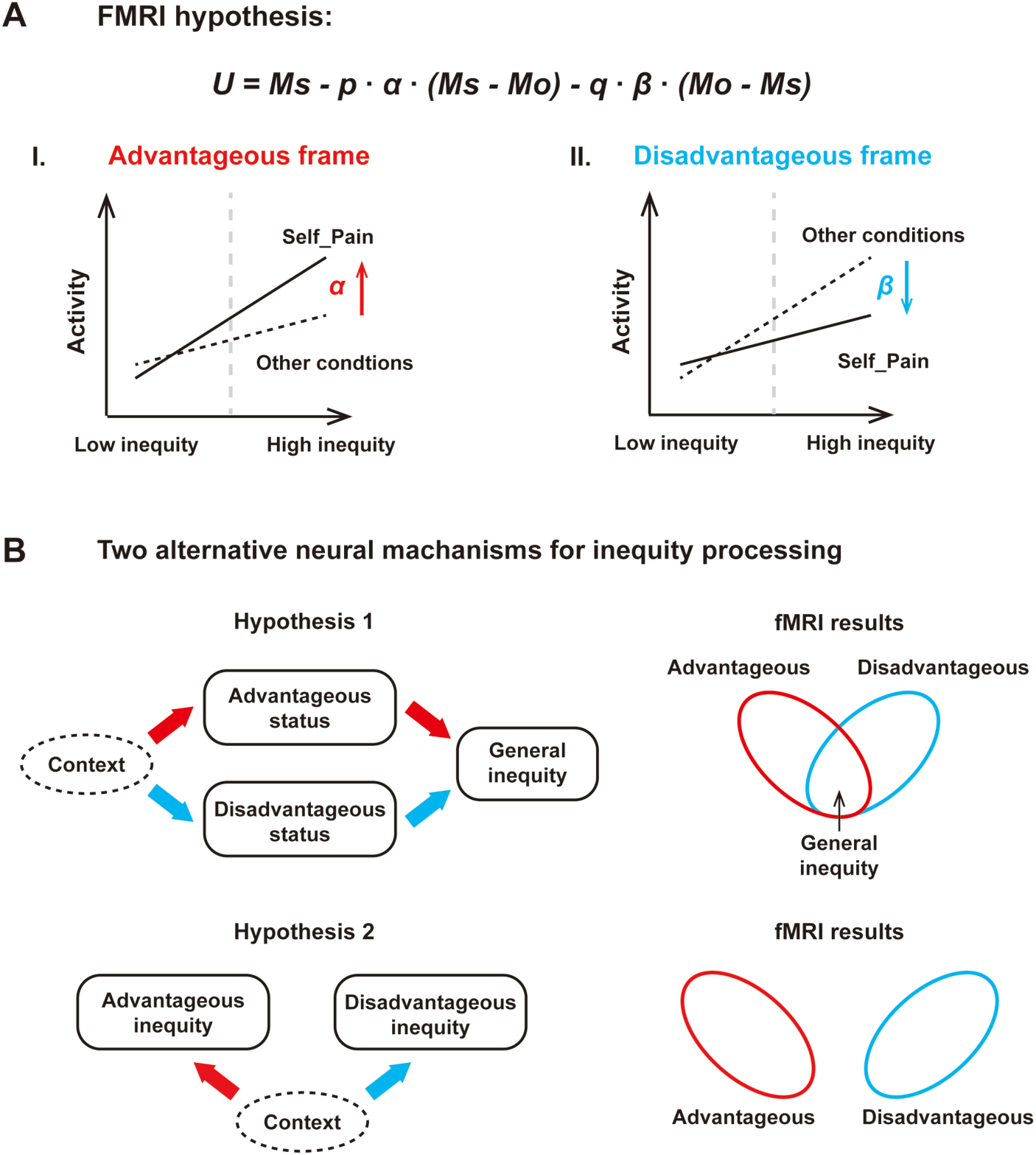
fMRI hypothesis. **(A)** In order to address the question as to whether there were dissociable brain systems for context-dependent advantageous inequity processing and disadvantageous inequity processing, binary choices were classified into an advantageous frame and a disadvantageous frame, and each type of choice was further medium split into high-inequity choices (HI) and low-inequity choices (LI). In close correspondence to the behavioral analysis, we established the fMRI hypotheses showing the potential response pattern for brain regions that were involved in context-dependent advantageous inequity and disadvantageous inequity aversion representation. **(A-I)** Given the increased advantageous inequity aversion induced by guilt, we hypothesized that brain regions representing context-dependent advantageous inequity aversion would show increased sensitivity to advantageous inequity in the Self_Pain condition, which would in turn result in boosted activity difference between HI and low LI conditions in these regions in the advantageous frame. Thus, the activity of these brain regions would show significant Agent × Outcome × Inequity level 3-way interaction effects in the advantageous frame. **(A-II)** Similarly, brain regions representing context-dependent disadvantageous inequity aversion would also show significant Agent × Outcome × Inequity level 3-way interaction effects, but the direction of this effect would be opposite to that the in advantageous frame. **(B)** Predications for fMRI results according to two alternative hypotheses regarding the neural mechanisms for inequity processing in the brain. According to the Hypothesis 1, if we can identify brain regions that show different sensitivity to inequity across contexts (i.e. the interaction effect between context and inequity level), we would get not only separate neural correlates for advantageous and disadvantageous status, but also observe overlapping brain regions, which represent general inequity **(B-I)**. According to Hypothesis 2 we should observe that the manipulation of context might modulates non-overlapping brain networks for advantageous and disadvantageous inequity **(B-II)**.

For the advantageous frame, to search for the neural correlates underlying context-dependent advantageous inequity aversion, we identified brain regions that showed significant Agent × Outcome × Inequity level 3-way interaction effects (voxel-level *P* < 0.001, cluster-level *P*_FWE_ < 0.05) (see *FMRI data analysis* for details). Significant activations were found in the left aINS [-30, 21, -20], rDLPFC [39, 20, 37], and DMPFC [-12, 47, 40] (Figure 4A, I and VI; Table S7). Compared with the other conditions, when the participant inflicted pain upon the co-player, the activity difference between HI and LI choices in left aINS, rDLPFC, and DMPFC increased in the advantageous frame (Figure 4A, II, IV and VII; Figure S3. A-C), indicating the increased sensitivity of these regions to advantageous inequity. The effects of left aINS, rDLPFC, and DMPFC were absent in disadvantageous frame (Figure 4A, III, V and VIII).

**Figure 4.**
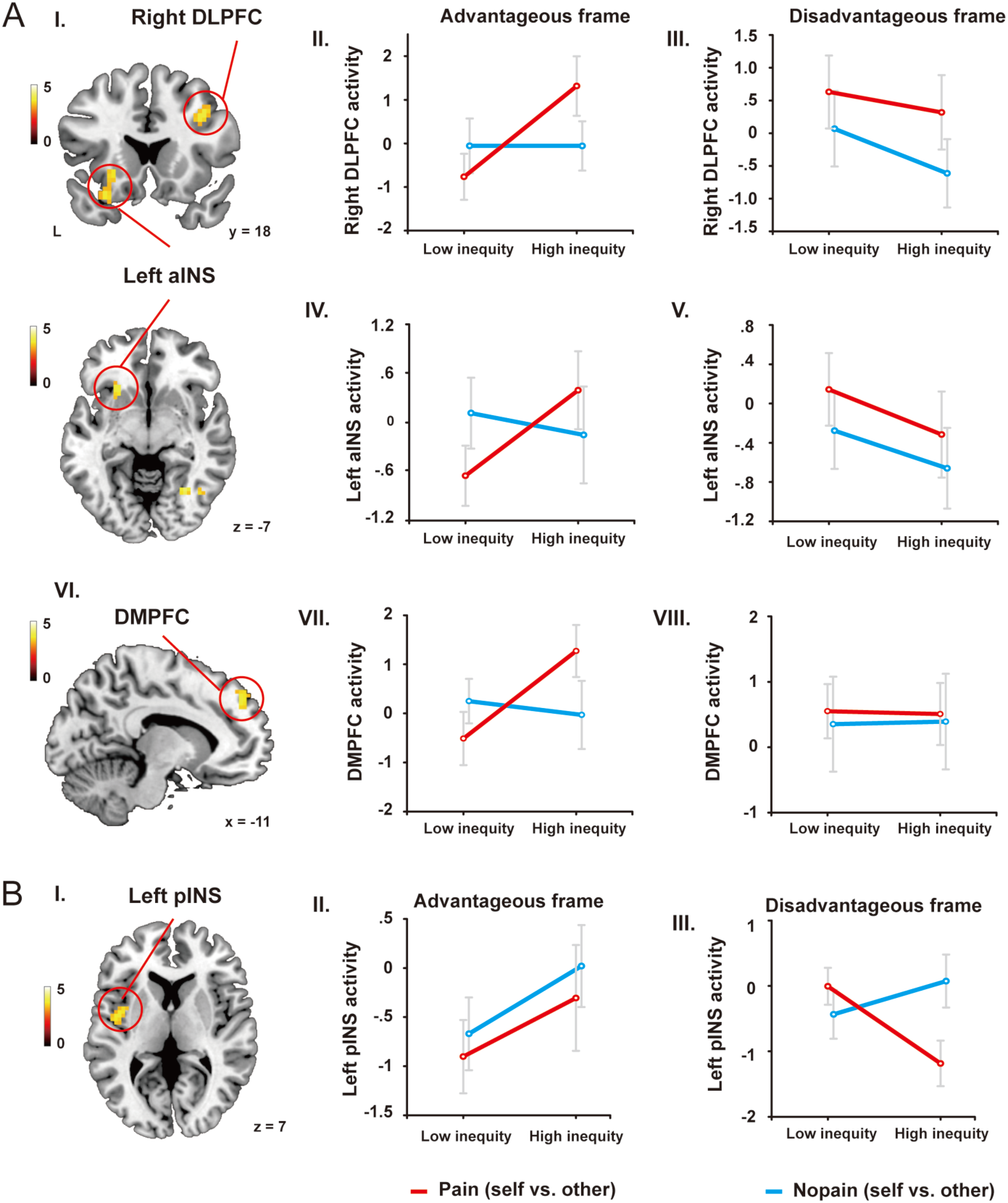
Neural representations of context-dependent advantageous inequity and disadvantageous inequity aversion. **(A)** Compared with the other conditions, when the participant inflicted pain upon the co-player (i.e. guilt condition), the activity difference between high inequity and low inequity choices in left aINS, rDLPFC, and DMPFC (I and VI) increased in the advantageous frame, indicating the increased sensitivity of these regions to advantageous inequity (II, IV, and VII). The effects of left aINS, rDLPFC, and DMPFC were absent in disadvantageous frame (III, V and VIII). **(B)** Compared with the other conditions, when the participant inflicted pain upon the co-player (i.e. guilt condition), the activity difference between high inequity and low inequity choices in left pINS (I) decreased in the disadvantageous frame, indicating the decreased sensitivity of these regions to disadvantageous inequity (III). The effect of left pINS was absent in the advantageous frame (II).

For the disadvantageous frame, we also identified brain regions that showed a significant between Agent × Outcome × Inequity level 3-way interaction effects, but the direction of this effect was opposite to that in advantageous frame (voxel-level *P* < 0.001, cluster-level *P*_FWE_ < 0.05) (see *FMRI data analysis* for details), as shown in the behavioral data. This contrast revealed a single activation in the left pINS [-42, -4, 7] (Figure 4B-I, Table S7), which showed reduced sensitivity to disadvantageous inequity in the Self_Pain condition, relative to the other conditions (Figure 4B-III; Figure S3D). The effect of left pINS was absent for the advantageous frame (Figure 4B-II). Further, no cluster survived from correction in either the conjunction analysis of the two frames or if we combined the data of the two frames to conduct contrast analysis. Our results demonstrated dissociable neural mechanism underlying context-dependent processing related to advantageous inequity aversion and disadvantageous inequity aversion: while left aINS, rDLPFC and DMPFC represent the context-dependent advantageous inequity aversion, left pINS represents the context-dependent disadvantageous inequity aversion.

### Functional Connectivity of the pINS in the Disadvantageous Frame

To test the probability that the processing of context-dependent advantageous and disadvantageous inequity aversion relied on not only on distinct neural activations but also distinct functional connectivity, we performed a psychophysiological interaction analysis (PPI; Friston et al., 1997) focusing on the left aINS, rDLPFC and DMPFC identified in the advantageous frame and the left pINS identified in the disadvantageous frame. When the left pINS was used as the seed, our results revealed significant Agent × Outcome × Inequity level 3-way interaction effects on the functional connectivity between left pINS and dACC ([-6, 20, 34]; Max *T*-value = 3.59; cluster size = 79; Figure 5A and B), and between left pINS and right amygdala ([27, 2, -20]; Max *T*-value = 3.26; cluster size = 22; Figure 5C and D) (SVC, *P*_FWE_ < 0.05, following an initial threshold of *P* < 0.005, uncorrected). Notably, the small volumes of dACC and amygdala were defined as spheres with 10 mm radius, centered on the peak MNI coordinates extracted from the meta-analyses on “emotion” and “conflict” terms in the Neurosynth database. Specifically, compared with Self_Nopain condition, the contrast between the HI condition and the LI condition in the left pINS-dACC connectivity and left pINS-right amygdala connectivity decreased in the Self_Pain condition (dACC: *F*_(1, 25)_ = 5.068, *p* = 0.033, right amygdala: *F*_(1, 25)_ =5.767, *p* = 0.024), while no difference was observed between the Other_Nopain condition and the Other_Pain condition (dACC: *F*_(1, 25)_ < 0.001, *p* = 0.983, right amygdala: *F*_(1, 25)_ = 0.275, *p* = 0.604). In contrast, PPI analyses with seeds identified in the advantageous frame failed to survive the whole-brain cluster level threshold and SVC.

**Figure 5.**
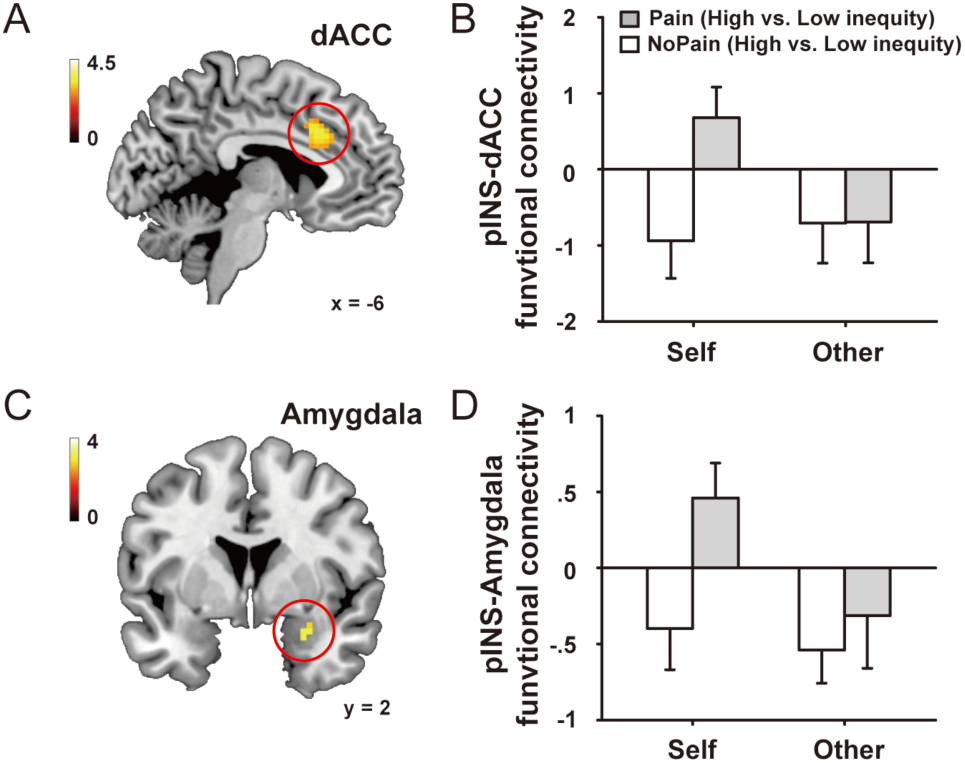
Functional connectivity of the pINS in disadvantageous frame. When the left pINS was used as the seed, our results revealed significant Agent × Outcome × Inequity level 3-way interaction effects on the functional connectivity between left pINS and dACC **(A)** and between left pINS and right amygdala **(C)** (SVC, *P*_FWE_ < 0.05, following an initial threshold of *P* < 0.005, uncorrected). Compared with Self_Nopain condition, the contrast between HI condition and LI condition for left pINS-dACC connectivity and left pINS-right amygdala connectivity decreased in Self_Pain condition (dACC: *F*_(1, 25)_ = 5.068, *p* = 0.033, right amygdala: *F*_(1, 25)_ =5.767, *p* = 0.024), while no difference was observed between Other_Nopain condition and Other_Pain condition (dACC: *F*_(1, 25)_ < 0.001, *p* = 0.983, right amygdala: *F*_(1, 25)_ = 0.275, *p* = 0.604) **(B and D)**. These results indicated that when participants felt guilty, left pINS-dACC connectivity and left pINS-right amygdala connectivity showed reduced representation of disadvantageous inequity, which was consistent with the decreased disadvantageous inequity aversion in this condition suggested by behavioral results.

### Spatial Gradient Within Insula for Context-dependent Advantageous Inequity vs. Disadvantageous Inequity Aversion Processing

For a more quantitative examination of this spatial segregation within the insula, we defined five regions of interest (ROIs) along the axis from the left aINS to the left pINS (Figure 6A), which were identified in the whole-brain analysis for the advantageous and disadvantageous frames. For each ROI, we extracted the beta estimates for each condition using fMRI data without smoothing in data preprocessing, evaluated the absolute Agent × Outcome × Inequity 3-way interaction effects for the advantageous frame and the disadvantageous frame. A 2 (context: Advantageous vs. Disadvantageous) × 5 (ROI locations) repeated-measures ANOVA showed a clear spatial distinction between the left aINS and the left pINS in the processing of context-dependent advantageous inequity vs. disadvantageous inequity aversion, respectively (interaction of context with ROI locations: *F*_(4, 100)_ = 4.319, *P* = 0.003; Figure 6B). The interaction effect for advantageous inequity was stronger in more anterior ROIs, whereas the interaction effect for disadvantageous inequity was stronger in more posterior ROIs. The difference in the linear effect of ROI location between contexts was also significant, *F*_(1, 25)_ = 7.583, *P* = 0.011. More specifically, the strength of the interaction effect for the disadvantageous frame increased linearly from the left aINS to the left pINS, *F*_(1, 25)_ = 5.084, *P* = 0.033, whereas the interaction effect for the advantageous frame showed the opposite trend, *F*_(1, 25)_ = 1.946, *P* = 0.175. The results remained the same using smoothed data (Figure S5).

**Figure 6.**
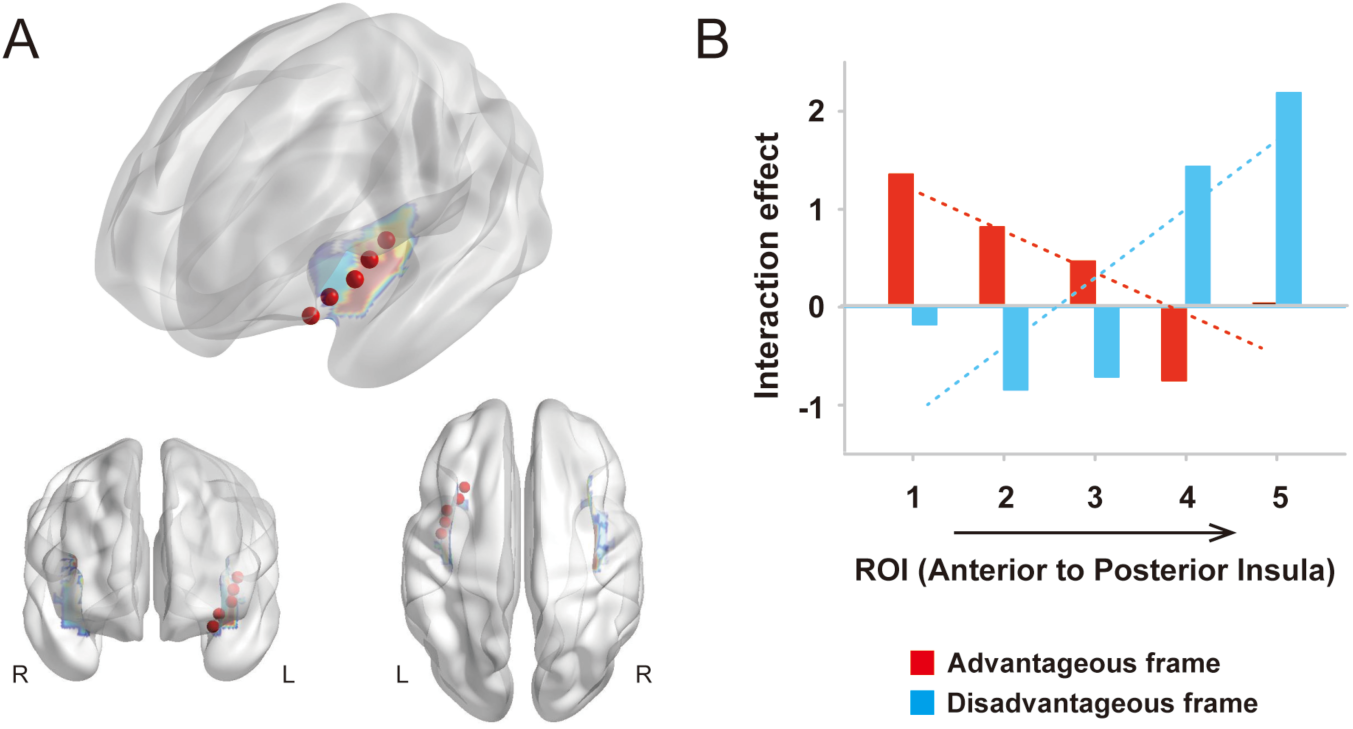
Spatial gradient for context-dependent inequity aversion representations. **(A)** Five regions of interest (ROIs) were defined along the axis from the left aINS to the left pINS, which were identified in the whole-brain analysis for the advantageous and disadvantageous frames. **(B)** For each ROI, we extracted the beta estimates for each condition using fMRI data without smoothing in data preprocessing, and evaluated the absolute 3-way interaction effects [(Self_Pain_High > Self_Pain_Low) − (Self_Nopain_High > Self_Nopain_Low) − (Other_Nopain_High > Other_Pain_Low) + (Other_Nopain_High > Other_Nopain_Low)] for advantageous frame and disadvantageous frame. The interaction effect for advantageous inequity was stronger in more anterior ROIs, whereas the interaction effect for disadvantageous inequity was stronger in more posterior ROIs. Dotted lines indicate linear fits of the spatial gradient for advantageous (red) and disadvantageous (blue) frames.

### Distinct Neural Mechanism Underlying Individual Differences in Contextual Effects on Advantageous and Disadvantageous Inequity Aversion

We further investigate whether neural mechanism underlying individual differences in the contextual effects on advantageous and disadvantageous inequity aversion were distinct from each other. Here, the strength of neural adjustments was defined as the value of 3-way interaction effects [(Self_Pain_High > Self_Pain_Low) − (Self_Nopain_High > Self_Nopain Low) − (Other_Nopain_High > Other_Pain_Low) + (Other_Nopain_High > Other_Nopain_Low)]; the more the difference between this value and zero, the stronger the neural adjustments are. Individual difference in sensitivity to context (i.e. guilt-proneness reflected by GASP guilt-NBE score) was related to the strength of neural adjustments by conducting ROI analyses (see *Methods* for details) for brain regions involved in advantageous inequity aversion processing (i.e. left aINS, rDLPFC, and DMPFC) and brain regions involved in disadvantageous inequity aversion processing (i.e. left pINS, ACC, and right amygdala). Results demonstrated that individual sensitivity to context predicted the strength of neural adjustments in rDLPFC (Figure 7A) in the advantageous frame (r = 0.437, *p* = 0.026), and the regression remained significant after excluding the extreme value, which could be considered an outlier (r = 0.402, *p* = 0.047) (Figure 7B, left). That is, the value of the interaction effect on rDLPFC activity in the advantageous frame was significantly above zero for participants with greater guilt proneness (N = 14), t_13_ = 2.899, *p* = 0.012, whereas this effect was absent for those with lower guilt proneness (N = 12), t_11_ = 1.124, *p* = 0.285 (Figure 7B, right). The association between individual proneness and rDLPFC activity was not observed for the disadvantageous frame (r = 0.018, *p* = 0.929). Moreover, individual guilt proneness could predict the strength of neural adjustments in dACC (Figure 7C) in disadvantageous frame (r = -0.404, *p* = 0.041) (Figure 7D, left). Compared with participants with low guilt proneness, participants with high guilt proneness showed significantly stronger neural adjustments in dACC (more negative interaction effect in [(Self_Pain_High > Self_Pain_Low) − (Self_Nopain_High > Self_Nopain_Low) − (Other_Nopain_High > Other_Pain_Low) + (Other_Nopain_High > Other_Nopain_Low)] contrast) in the disadvantageous frame, t_24_ = 2.472, *p* = 0.021 (Figure 7D, right). This relationship between individual proneness and dACC activity was not observed for the advantageous frame (r = 0.129, *p* = 0.529).

**Figure 7.**
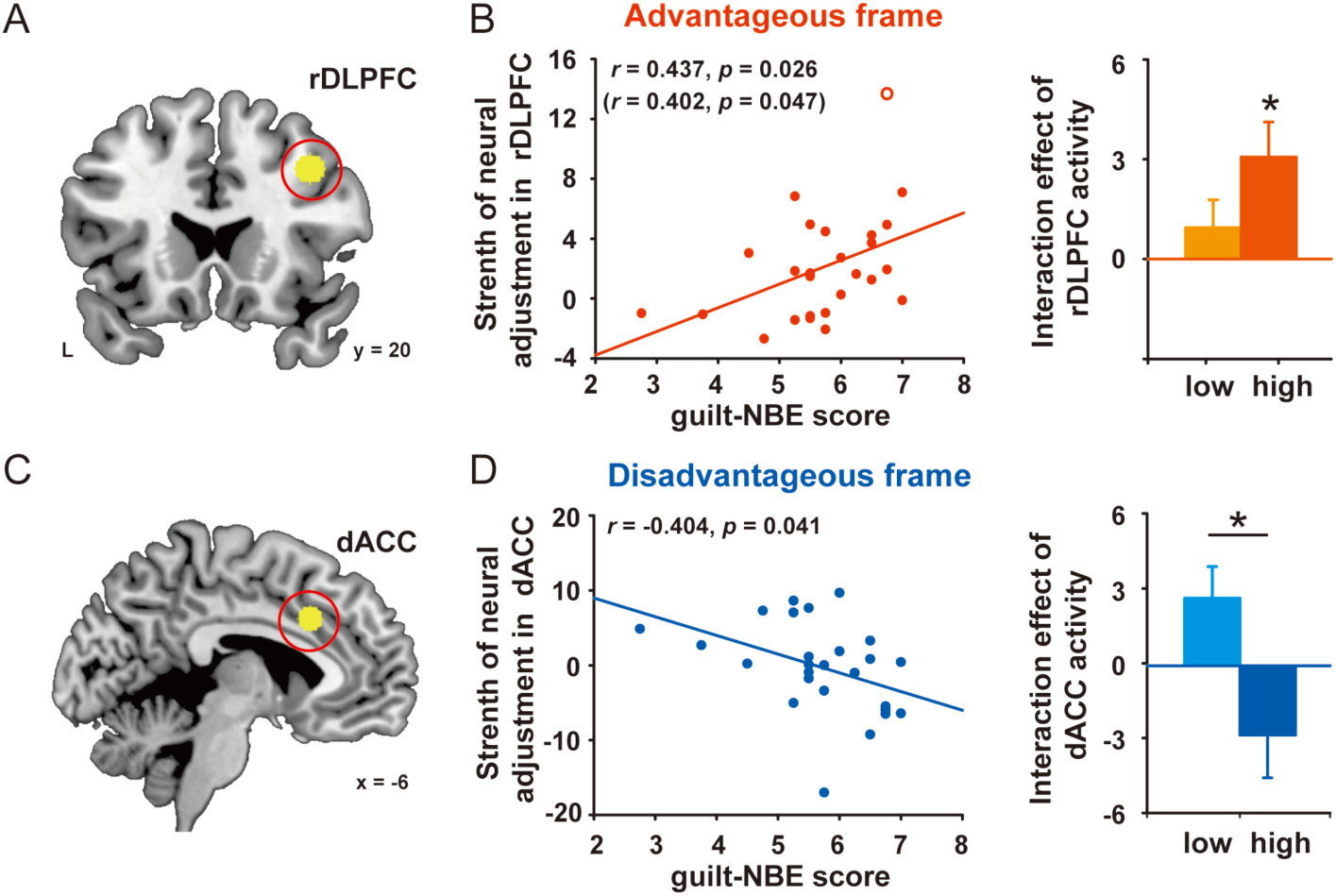
Dissociable neural mechanism underlying individual differences in contextual effects on advantageous and disadvantageous inequity aversion. Individual guilt proneness (i.e. guilt-NBE score) predicted the strength of neural adjustments in rDLPFC in the advantageous frame *(r* = 0.437, *p* = 0.026), and the regression remained significant after excluding the extreme value, which could be considered an outlier (r = 0.402, *p* = 0.047) (A and B, left). The value of the interaction effect on rDLPFC activity in the advantageous frame was significantly above zero for participants with greater guilt proneness (N = 14), *t*_13_ = 2.899, *p* = 0.012, whereas this effect was absent for those with lower guilt proneness (N = 12), *t*_11_ = 1.124, *p* = 0.285 (B, right). The association between individual proneness and rDLPFC activity was not observed for the disadvantageous frame (*r* = 0.018, *p* = 0.929). Moreover, individual guilt proneness was predictable of the strength of neural adjustments in dACC in disadvantageous frame (*r* = -0.404, *p* = 0.041) (C and D, left). Compared with participants with low guilt proneness, participants with high guilt proneness showed significantly stronger neural adjustments in dACC (more negative interaction effect) in the disadvantageous frame, *t*_24_ = 2.472, *p* = 0.021 (D, right). This relationship between individual guilt proneness and dACC activity was not observed for advantageous frame (r = 0.129, *p* = 0.529).

We established two more GLMs to exclude the possible confounding of the value representation of self-payoff (GLM2) and other-payoff (GLM3). For the fMRI analyses on the effect of guilt on self-payoff and other-payoff representation, no cluster survived the whole-brain cluster level threshold. To further ensure that the specific role of left aINS, rDLPFC, and DMPFC in advantageous inequity aversion processing, and the specific role of left pINS in disadvantageous inequity aversion processing, we extracted and plotted the beta estimates of these 4 regions for 8 conditions for self-payoff (i.e. Self_Pain_LS, Self_Pain_HS, Self_Nopain_LS, Self_Nopain_HS, Other_Pain_LS, Other_Nopain_HS, Other_Nopain_LS, and Other_Nopain_HS) in Figure S4, A-D. Similarly, we extracted and plotted the beta estimates of these 4 regions for 8 conditions for other-payoff (i.e. Self_Pain_LO, Self_Pain_HO, Self_Nopain_LO, Self_Nopain_HO, Other_Pain_LO, Other_Nopain_HO, Other_Nopain_LO, and Other_Nopain_HO) in Figure S4, E-H. No 3-way interaction effect was found for neural responses to self-payoffs or other-payoffs in left aINS, rDLPFC, DMPFC, and left pINS, indicating the observed effects of these regions in inequity processing were not driven by the confounding value representations of self-payoffs or other-payoffs.

## Discussion

Combining an interpersonal interactive game (Yu et al., 2014a) that modulated individual advantageous and disadvantageous inequity aversion simultaneously and a variant of DG (Saez et al., 2015) that enabled us to characterize individual changes in inequity aversion, our results provide advanced evidence demonstrating two distinct types of social context-dependent inequity aversion processing mechanisms in the human brain, such that the processing of advantageous inequity aversion involves the left aINS, the rDLPFC, and DMPFC, while the processing of disadvantageous inequity aversion involves the left pINS, the amygdala and dACC. These results are consistent with previous evidence from of behavioral economics (Dawes et al., 2007; Fehr and Schmidt, 1999; Loewenstein et al., 1989; Van den Bos et al., 2006) and from developmental (Blake et al., 2015; Blake and McAuliffe, 2011; Fehr et al., 2008; McAuliffe et al., 2013; McAuliffe et al., 2017), and evolutionary (Brosnan, 2009; Brosnan and de Waal, 2014) perspectives.

### Distinct roles of insular sub-regions in inequity aversion processing

The insula is involved in a circuit responsible for the detection of salience (for a review, see Menon and Uddin, 2010), which integrates external elements about stimuli and internal information about individual cognitive, homeostatic, and emotional states to organize behavior. Previous studies employing a diverse range of methodological approaches—including cytoarchitectonic mapping (Kurth et al., 2010a; Mesulam and Mufson, 1982), tractography (Cerliani et al., 2012; Nanetti et al., 2009), meta-analysis of task-related fMRI data (Kurth et al., 2010b; Mutschler et al., 2009; Wager and Barrett, 2004), and functional connectivity (Cauda et al., 2011; Deen et al., 2011; Nelson et al., 2010)—converge on the existence of an antero-posterior organization of the insula, in which the posterior region is involved in primary interoceptive representation, whereas the anterior region is involved in motivational, social, and cognitive processing (Craig, 2009). Extending this functional organization of the insula, our results show that the context-dependent representation of inequity aversion exhibited a spatial gradient in activity within the insula, such that anterior parts are predominantly involved in advantageous inequity aversion and posterior parts are predominantly involved in disadvantageous inequity aversion.

The involvement of pINS in disadvantageous inequity aversion processing in the current study might seem puzzling, considering the primary role of pINS in interoceptive representations (Craig, 2009) and the role of aINS in UG responders’ responses to unfair (i.e. disadvantageous) offers (For reviews and meta-analyses, see Aoki et al., 2015; Fehr and Camerer, 2007; Feng et al., 2015; Gabay et al., 2014). However, by mapping peak coordinates of insula regions identified in studies focused on UG responders on insular sub-regions, we found that no matter what the context is, unfair offers were associated with increased activity in the dorsal anterior insula, whereas the context-related processing of unfair offers involved both the dorsal anterior and the mid-posterior parts of insula, which is actually consistent with the pINS identified here. The involvement of pINS in inequity processing is further supported by Hsu et al. (2008), who demonstrated that pINS represented the aversion to inequity in a third-party resource distribution task. These results are also in line with the involvement of pINS in other high-level computations related to inter-temporal choice (Tanaka et al., 2004; Wittmann et al., 2007) and language perception (Jones et al., 2010).

Here we did not observe the role of aINS in context-dependent disadvantageous inequity processing as suggested by previous UG studies (For reviews and meta-analyses, see Aoki et al., 2015; Fehr and Camerer, 2007; Feng et al., 2015; Gabay et al., 2014). One possible explanation is that, in contrast to the passive responses to unfair offers in UG, here we focused on the decision-making process when individuals made monetary allocation voluntarily as a dictator. Since the payoffs of oneself and the other were determined by participants’ decisions, they did not need to detect the fairness norm violation by others, which is suggested to be the main role of aINS in UG (Chang and Sanfey, 2013; Cheng et al., 2015; Civai et al., 2012; Corradi-Dell'Acqua et al., 2012; Gu et al., 2015; Guo et al., 2013; Xiang et al., 2013; Zheng et al., 2014). Moreover, it is shown that the experience of negative emotions elicited by receiving less than someone else, which is related to the anticipated experience of “envy” (Fehr and Schmidt, 1999; Jankowski and Takahashi, 2014; Krajbich et al., 2009; Ortony et al., 1988; Rey - Biel, 2008; Steinbeis and Singer, 2013), is a key component for the emergence of disadvantageous inequity aversion (McAuliffe et al., 2017). For example, in a reward distribution task, the activity in amygdala, a region that is implicated in negative emotions (Barrett et al., 2007; Feinstein et al., 2011; Phelps, 2006), correlated positively with subjects’ willingness to reduce others’ outcomes to create equality (i.e. reducing disadvantageous inequity; Yu et al., 2014b). The activity in amygdala was also suggested to reflect the negative emotional responses to disadvantageous offers for UG responders (Gospic et al., 2011; Haruno and Frith, 2010; Haruno et al., 2014). Notably, previous studies have shown both structural (McDonald, 1998) and functional (Roy et al., 2009) connections between pINS and amygdala. Patients with the PTSD symptom of hyperarousal are shown to have hyperconnectivity between the pINS and amygdala, thus indicating the involvement of pINS-amygdala connectivity in emotional response (Sripada et al., 2012). One previous study (Dunn et al., 2012) has shown that interoceptive awareness, which is mainly represented in pINS (Craig, 2011; Craig, 2009), moderates the emotional responses to unfair proposals in UG. Here, we consistently found here that not only pINS activity but also its functional connectivity with amygdala showed a significant interaction effect between context and disadvantageous inequity aversion level. Based on these results, we suggest that the adjustment of disadvantageous inequity aversion according to the social contexts in DG involved the neural adjustments in the interoceptive system and its interaction with emotional systems, which might modulate the emotional responses to disadvantageous options.

In contrast to the absent effect of aINS in the disadvantageous frame, we observed the context-dependent response in aINS in advantageous frame. In DG, the advantageous inequity aversion is usually explained as the anticipated “guilt” feeling, i.e. the negative feeling induced by norm violation (i.e. earning more than others; Fehr and Schmidt, 1999; Krajbich et al., 2009; Rey - Biel, 2008). Chang et al. (2011) investigated the role of anticipated guilt in motivating cooperative behaviors while participants decided whether or not to return some money to a co-player who had invested money in them (i.e. the trust game). Their neuroimaging results indicated the role of aINS in minimizing anticipated guilt and motivating adherence to the perceived social norm derived from the predicted others’ expectations. Consistently, previous work on decisions to conform to a perceived social norm has revealed the involvement of aINS (Berns et al., 2010; Klucharev et al., 2009), which indicates that the function of aINS is to track deviations from the perceived social norm and bias actions to maintain adherence to this norm. This opinion can also be applied to a series of studies demonstrating the involvement of aINS in the decision to reject unfair offers in UG (Chang and Sanfey, 2013; Cheng et al., 2015; Civai et al., 2012; Corradi-Dell'Acqua et al., 2012; Gu et al., 2015; Guo et al., 2013; Sanfey et al., 2003; Xiang et al., 2013; Zheng et al., 2014). In the current study, our results extended the role of aINS to support individual adjustment of advantageous inequity aversion (or the anticipated guilt) according to social contexts. In light of the function of aINS suggested by the above studies, we suggested that the increased sensitivity of aINS responses to advantageous inequity when participants inflicted pain upon others might reflect their increased sensitivity to anticipated norm violation in choosing advantageous option, which in turn lead to behaviorally increased aversion to advantageous inequity.

### Neural correlates of context-dependent advantageous inequity aversion

In addition to aINS, two regions that play critical roles in social decision-making, rDLPFC and DMPFC, were involved in the processing of context-dependent advantageous inequity aversion. DMPFC is primarily related to the understanding of other's mental states (i.e. mentalizing; For review, see Frith and Frith, 2003; Isoda and Noritake, 2013; Lieberman, 2007), with strong anatomical and functional connectivity to other mentalizing network nodes, including the temporo-parietal junction (Buckner et al., 2008) and the DLPFC (Duncan, 2010). Notably, the ability of mentalizing (i.e. understanding the other’s feeling of being hurt) is the foundation of guilt experience (Basil et al., 2008; Baumeister et al., 1994). Thus, it is reasonable to further extend the mentalizing process to the processing of the anticipated “guilt” feeling, i.e. the advantageous inequity aversion (Fehr and Schmidt, 1999; Krajbich et al., 2009; Rey - Biel, 2008). The recruitment of DMPFC in the context-dependent processing of advantageous inequity here may help individuals to anticipate the co-player's feelings of disappointment in response to getting less in different contexts more accurately and lead to downstream behavioral adjustment in accordance with the context.

Previous studies suggest that disadvantageous inequity aversion emerges in early childhood, whereas advantageous inequity aversion emerges in late childhood, as it may require the development of behavioral-control-related brain regions to support norm compliance (For review, see McAuliffe et al., 2017). Consistently, a behavioral study demonstrated that rejecting advantageous inequity to themselves requires more cognitive resources than rejecting disadvantageous inequity to themselves (Van den Bos et al., 2006). Here, we provided neural evidence that DLPFC, a region implicated in cognitive control (Buckholtz and Marois, 2012; Miller and Cohen, 2001) and social norm compliance (Knoch et al., 2006; Nihonsugi et al., 2015; Ruff et al., 2013; Zhu et al., 2014), contributed to the adjustment of advantageous inequity aversion to social contexts, but not in the disadvantageous frame. Moreover, individuals with greater interaction effects between context and advantageous inequity in DLPFC activity here was associated with higher sensitivity to guilt context in daily life; this finding is congruent with the suggestion that robust cognitive control allows for responding to the dynamic environments with increased flexibility (Steinbeis and Crone, 2016). Taken together, our findings indicated the “social” nature of advantageous inequity aversion, which may recruit both the cognitive control process represented in the DLPFC and the mentalizing process represented in the DMPFC to adjust the signal of perceived social norm violation from aINS according to the contexts.

### Neural correlates of context-dependent disadvantageous inequity aversion

In addition to the pINS-amygdala functional connectivity discussed above, context-dependent disadvantageous inequity aversion processing was also associated with the functional connectivity between pINS and dACC, a region implicated in conflict monitoring (For review, Heilbronner and Hayden, 2016). Previous studies have shown increased activity of dACC in response to unfair offers in UG (Aoki et al., 2015; Fehr and Camerer, 2007; Feng et al., 2015; Gabay et al., 2014), which may reflect the conflict between the unfairness-evoked aversive responses and the self-interest to gain monetary reward (Fehr and Camerer, 2007; Sanfey et al., 2003). Thus, we speculated that, during the experience of guilt, the decreased sensitivity of pINS-dACC functional connectivity to disadvantageous inequity might indicate reduced conflict between aversive responses and self-interest in this condition. The role of dACC in context-dependent disadvantageous inequity aversion was further confirmed by the correlation between individual sensitivity to contexts and the strength of neural adjustment in dACC. In sum of our findings in disadvantageous frame, the context-dependent processing of disadvantageous inequity aversion may depend on the neural adjustments in the interoceptive system, emotional system and conflict monitoring system to adjust the negative emotional responses to disadvantageous inequity and its conflict with self-interest.

### Implications for future studies

First, in the current study we combined interactive games in social psychology and computational models in neuroeconomics to provide the first evidence for the neural mechanisms underlying context-dependent processing of advantageous and disadvantageous inequity. While interactive games enabled us to observe participants’ behaviors and neural responses in real life-like contexts, the application of sophisticated economic models enabled us to quantify psychological constructs mathematically and then examine these psychological constructs at neural level (Crockett, 2016; O'doherty et al., 2007). Although in the current study we can only provide evidence for a single context (i.e. the guilt context), our interdisciplinary paradigm may promote future studies to investigate this question in various contexts. Second, our findings provide valuable evidence to help us distinguish between two alternative hypotheses regarding the neural mechanisms for inequity processing from the view of context-dependency; this ability provides us with several advantages compared with previous studies on the two forms of inequity aversion in decontextualized environments (Fliessbach et al., 2012; Güroğlu et al., 2014; Yu et al., 2014b). Although these previous studies extended our understanding on the neural bases underlying two forms of inequity aversion by comparing participants’ neural responses in unequal conditions with those in equal conditions, one problem with this kind of analysis is the inherent correlation between self-payoff and advantageous inequity and the correlation between other-payoff and disadvantageous inequity. Thus the observed neural results may also reflect the representations of the self-payoff or the other-payoff and not just the representations of advantageous inequity or disadvantageous inequity, which may account for the relative inconsistent neural results of these previous studies. By comparing neural responses to the same unequal options under different contexts, our experiments to some extent exclude the confounding of self-payoffs and other-payoffs. Our findings are waiting future independent replications. Future studies using multi-voxel level analysis (e.g. MVPA or RSA) and causal manipulations (e.g. TMS) may provide more evidence for these questions. Third, the ability to flexibly integrate contextual information and adapt decisions and behavior accordingly is a crucial skill underlying successful social interaction (i.e. social context sensitivity; Obradović and Boyce, 2009; Steinbeis and Crone, 2016). Understanding how individual make behavioral and neural adjustments to the social context may provide valuable insight regarding certain social dysfunctions, such as autism (Palmer et al., 2015) or psychopathy (Domes et al., 2013), which are associated with reduced sensitivity to social signals. Our results suggested that the strength of neural adjustments in rDLPFC in the advantageous frame and dACC in the disadvantageous frame were positively correlated with individual sensitivity to guilt context in daily life. Future work may attempt to test whether this individual difference in neural adjustments can be applied to other social contexts that influence inequity aversion and research on social dysfunctions.

In conclusion, the current study illustrates the dissociable neural mechanisms underlying context-dependent processing of advantageous and disadvantageous inequity aversion. These results shed light on how individuals integrate social context information into the decision-making process and adjust behaviors accordingly and provide implications for research on dysfunctions involving social maladaptation such as autism and psychopathy.

## Acknowledgements

This work was supported by National Basic Research Program of China (973 Program: 2015CB856400) and National Natural Science Foundation of China (91232708, 31170972, 31630034). The behavioral part of this study has been presented as a poster at the annual meeting of the Social and Affective Neuroscience Society 2016 (New York City, April 28-May 2, 2016). The whole study has been presented as a talk at the Society for Neuroeconomics Annual Conference 2017 (Toronto, October 6–8, 2017). The authors thank Dr. Christian C. Ruff, Dr. Peter Sokol-Hessner and Dr. Luke J. Chang for their advice on this paper.

## Footnotes

1 In the dictator game (DG), the participant determines how to split an endowment between himself/herself and a recipient as a "the dictator". The recipient simply receives the endowment given by the dictator and has no right to influence the outcome of the game.

2 In the ultimatum game (UG), the proposer receives a sum of money and proposes how to divide the sum between himself/herself and the responder. The responder chooses to either accept or reject this proposal. If the responder accepts, the money is split according to the proposal. If the responder rejects, neither the proposer nor the responder receives any money.

